# Optimal structure of metaplasticity for adaptive learning

**DOI:** 10.1101/129619

**Authors:** Peyman Khorsand, Alireza Soltani

## Abstract

Learning from reward feedback in a changing environment requires a high degree of adaptability, yet the precise estimation of reward information demands slow updates. We show that this tradeoff between adaptability and precision, which is present in standard reinforcement-learning models, can be substantially overcome via reward-dependent metaplasticity (reward-dependent synaptic changes that do not always alter synaptic efficacy). Metaplastic synapses achieve both adaptability and precision by forming two separate sets of meta-states: reservoirs and buffers. Synapses in reservoir meta-states do not change their efficacy upon reward feedback, whereas those in buffer meta-states can change their efficacy. Rapid changes in efficacy are limited to synapses occupying buffers, creating a bottleneck that reduces noise without significantly decreasing adaptability. In contrast, more-populated reservoirs can generate a strong signal without manifesting any observable plasticity. We suggest that ubiquitous unreliability of synaptic changes evinces metaplasticity that can provide a robust mechanism for adaptive learning.

## Introduction

To successfully learn from reward feedback, the brain must adjust how it responds to and integrates reward outcomes, since reward contingencies can unpredictably change over time (Behrens et al., 2007; Rushworth and Behrens, 2008). At the heart of this learning problem is a tradeoff between adaptability and precision. On one hand, the brain must rapidly update reward values in response to changes in the environment, and on the other hand, in the absence of any such changes, it must obtain accurate estimates of those values. This tradeoff, which we refer to as the adaptability−precision tradeoff (Farashahi et al., 2017a,b; Khorsand, Farashahi, and Soltani, 2016, Society for Neuroscience abstract), can be easily demonstrated in the framework of reinforcement learning (Sutton and Barto, 1998). According to this framework, larger learning rates result in higher adaptability but lower precision, and smaller learning rates give rise to lower adaptability but higher precision (Supplementary Note 1 and Supplementary Figure 1). In recent years, the failure of conventional reinforcement learning (RL) models to capture the level of adaptability and precision demonstrated by humans and animals has led to alternative explanations for how we deal with uncertainty and volatility in the environment (Behrens et al., 2007; Krugel et al., 2009; Payzan-LeNesture et al., 2011; Costa et al., 2015). However, most of these solutions for adjusting learning require complicated calculations, and their underlying neural substrates are unknown.

Given the central role of synapses in learning, we asked whether there are synaptic mechanisms that can adjust the brain’s level of plasticity according to reward statistics and, therefore, allow the learning process to be adaptable. A candidate mechanism for such adjustment is metaplasticity, defined as changes in the synaptic state that shape the direction, magnitude, and duration of future synaptic changes without any observable change in the efficacy of synaptic transmission (Abraham and Bear, 1996; Abraham 2008; Fusi et al., 2005; Muller-Dahlhaus and Ziemann, 2015; Yger and Gilson, 2015). Extending our recent heuristic model of reward-dependent metaplasticity, which exhibits the adjustment of learning to reward uncertainty (Farashahi et al., 2017a), here we examined a general class of metaplastic models to identify features that are beneficial for adaptive learning and for mitigating the adaptability−precision tradeoff (APT).

We found that the APT can be substantially overcome in superior metaplastic models. These models achieve both adaptability and precision by forming two separate sets of meta-states: reservoirs and buffers. Synapses in reservoir meta-states do not change their efficacy upon reward feedback whereas those in buffer meta-states can change their efficacy. In superior models, rapid changes in efficacy are limited to synapses occupying buffers, creating a bottleneck that reduces noise without significantly decreasing adaptability. In contrast, more-populated reservoirs can generate a strong signal without manifesting any observable plasticity. Finally, we showed that metaplastic transitions are crucial for adaptive learning since replacing these transitions with plastic ones reduce the ability of the model in mitigating the APT. Altogether, our results illustrate how metaplasticity can mitigate one of the most fundamental tradeoff in learning and moreover, reveal the critical features of metaplasticity that contribute to adaptive learning.

## Results

To study the relationship between adaptability and precision, we considered a general problem of estimating reward probability (*p*_*r*_) from a stream of binary outcomes (reward, no reward). Our general model of metaplasticity consisted of multiple meta-states associated with one of the two values of synaptic efficacy (weak and strong), and all possible transitions between these meta-states (Fig. 1a; see Methods). The difference between the fractions of synapses that are in the strong and weak meta-states determines the signal (*S*) stored in these synapses. Importantly, we assumed that metaplastic transitions have a consistent order, and thus, within the set of weak and strong meta-states, there are multiple meta-states with different levels of depth (Fig. 1a).

**Figure 1.**
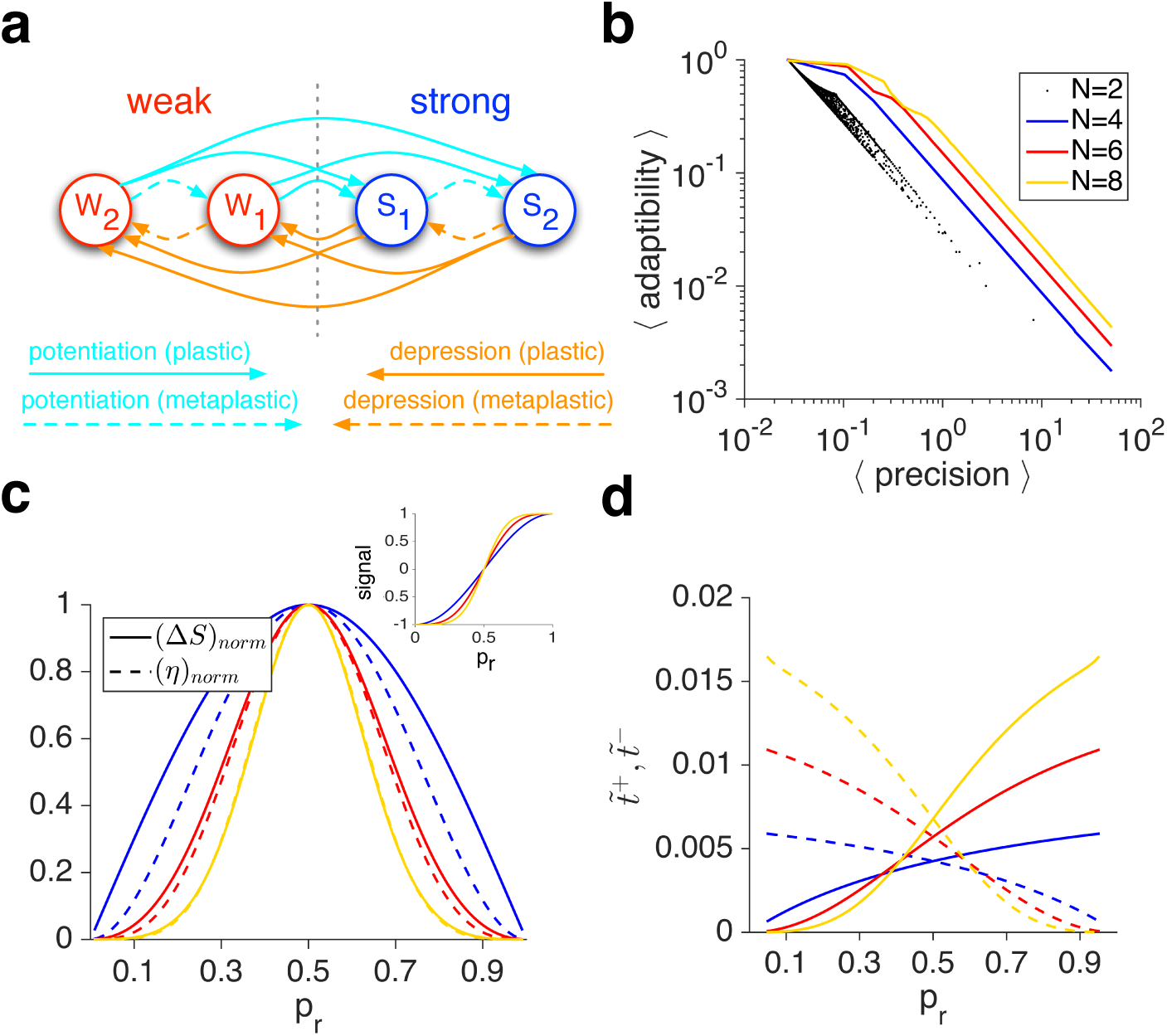
A general model of metaplasticity with ordered meta-states, and its behavior. (**a**) A schematic of the model. Synapses can transition between different meta-states with either weak or strong synaptic efficacies. (**b**) The APT in the plastic model (*N* = 2), and superior metaplastic models with different numbers of meta-states. Plotted is the average adaptability (over different values of *p*_*r*_) as a function of the average precision in different models. For the plastic model, each dot corresponds to a specific set of parameter values. For metaplastic models (*N* = 4, 6, 8), each outline connects models with optimized adaptability for a given value of average precision. (**c**) Matching of the sensitivity to noise in the metaplastic models. Plotted are the normalized sensitivity (Δ*S*) and one-step noise (*η*) as a function of *p*_*r*_ for three examples of superior metaplastic models with different numbers of meta-states. The sensitivity profile better matches the noise profile as *N* increases. (**d**) Adjustment of learning to reward probability in metaplastic models. Plotted are the effective learning rates for potentiation (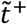, solid curves) and depression (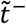, dashed curves) events as a function of *p*_*r*_. The effective learning rate on potentiation (depression) events increases (decreases) as reward probability increases, with a crossover at *p*_*r*_ = 0.5.

For a given value of reward probability, *p*_*r*_, the steady state of the model can be used to calculate the signal, and the weighted average change in signal due to single potentiation and depression events (‘one-step’ noise) provides a good proxy for noise (see Methods). To quantify how unambiguously the signal in a model differentiates between adjacent reward probabilities, we defined a quantity, termed the ‘precision’ (𝕡), equal to the sensitivity of the model’s signal to changes in reward probabilities (‘sensitivity’), divided by noise in the signal. Finally, the ‘adaptability’ (𝔸) of a model in estimating reward probability was defined as the rate at which the signal approaches its final value (see Methods). Since we were interested in conditions under which metaplasticity can improve the APT, we examined ‘superior’ metaplastic models (i.e. those which optimized 𝔸× 𝕡 for a given value of 𝕡).

We found that for many model parameters, the APT can be mitigated by superior metaplastic models that consist of as few as four meta-states (Fig. 1b; Supplementary Figure 2). These superior models overcame the APT by exhibiting two important characteristics, matching the sensitivity to noise and optimal adaptability. Firstly, the sensitivity of the signal to the reward probability matched the level of noise (sensitivity-to-noise matching), and this matching was improved with larger numbers of meta-states (Fig. 1c). Secondly, the adaptability of the models was optimized for a given level of noise (see below).

The first characteristic of superior models, the match between the sensitivity and the noise level, occurred because learning was naturally adjusted according to reward probability without any changes in the model’s parameters. To show this adjustment, we computed the ‘effective’ learning rates for potentiation and depression events for a given value of *p*_*r*_ (see Methods). The effective learning rate assigned a single rate to transitions between the weak and strong meta-states or vice versa (plastic transitions, Fig. 1a), which are the only transitions that can change synaptic efficacy and thus the signal. We found that the effective learning rate on rewarded trials 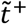 was very small for small values of *p*_*r*_ but monotonically increased as *p*_*r*_ increased (Fig. 1d). At the same time, the effective learning rate on unrewarded trials 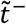 was large when *p*_*r*_ was close to zero and decreased as *p*_*r*_ increased. The effective learning rates on rewarded and unrewarded trials crossed over at 0.5

These complementary adjustments in learning resulted in a sigmoid-shape signal for superior, metaplastic models (Fig. 1c inset). Therefore, the maximum sensitivity (Δ*S*/Δ*p*_*r*_) for superior models occurred at *p*_*r*_ = 0.5, such that the steepest part of the signal matched the maximum level of noise. Importantly, the slope of the signal (i.e. sensitivity) at *p*_*r*_ = 0.5 was linearly proportional to the ratio of effective learning rates around *p*_*r*_ = 0.5 (results not shown), indicative of a direct relationship between the sensitivity-to-noise matching and adjustment of learning to reward probability. This outcome occurs in metaplastic models, without any changes in parameters, because as reward probability deviates from 0.5 (say when *p*_*r*_ > 0.5), more synapses move to shallower weak meta-states, increasing the effective potentiation rate above the effective depression rate (Fig. 1d). As the ratio of effective potentiation to depression rates increases, however, the fraction of synapses in weak meta-states decreases. Consequently, sensitivity to reward probability decreases as *p*_*r*_ becomes larger or smaller than 0.5. Finally, for a given level of precision, the signal became a steeper function of reward probability (maximum sensitivity increased) as the number of meta-states increased (Fig. 1c inset).

As noted above, another characteristic of superior models, in addition to the sensitivity-to-noise matching, was that their adaptability was optimized for a given level of noise. This optimization occurred because metaplasticity enabled superior models to form two separate sets of meta-states: reservoirs and buffers. Reservoirs, which are unique to metaplastic models, are the deepest sets of meta-states that cannot change their efficacy upon potentiation or depression events; they can only undergo metaplastic transitions (Fig. 2a). Buffers, on the other hand, are the shallowest meta-states, and are able to undergo plastic transitions that change their synaptic efficacy. The remainder of the meta-states provide transient meta-states. Because the superior models had reservoirs and buffers, they were able to keep a large proportion of their synapses in the weak or strong reservoirs (Fig. 2b). Synapses within reservoirs were protected against changes in efficacy upon potentiation or depression events, and as a result, the signal could increase without increasing the level of noise.

**Figure 2.**
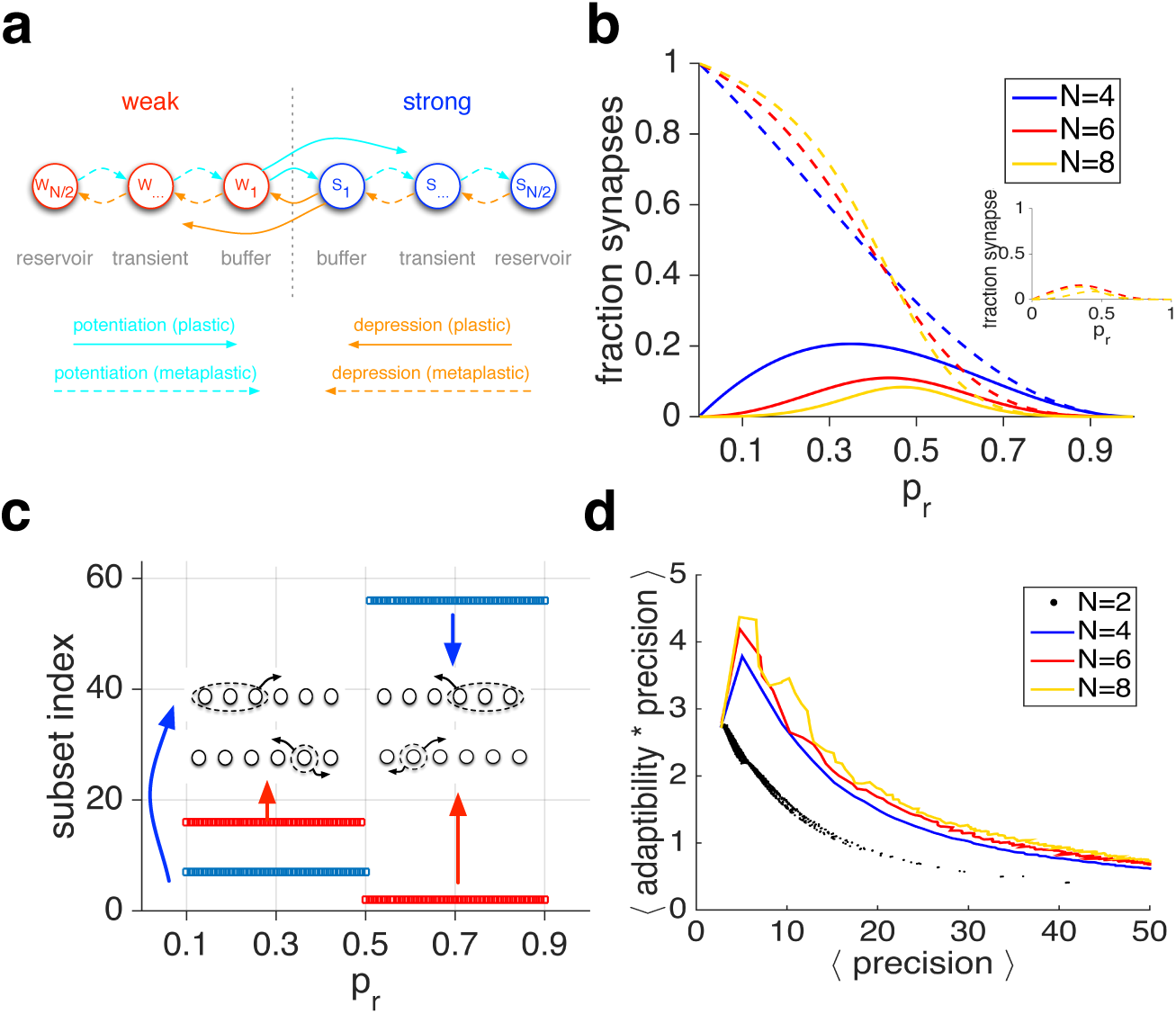
(**a**) Schematic of the reservoirs, buffers, and transient meta-states in metaplastic models. (**b**) Fractions of synapses in the weak reservoir (dashed lines), buffer (solid lines), and transient (inset) meta-states as a function of reward probability. Fractions in the strong reservoir, buffer, and transient meta-states are the mirror image (along *p*_*r*_ = 0.5) of their weak counterparts. As reward probability approaches 0.5, more synapses occupy transient and buffer meta-states, making the model more adaptable. As *p*_*r*_ deviates from 0.5, more synapses transition to reservoirs, enabling the model to protect the signal. (**c**) The subset of meta-states with the fastest and slowest effective transition rates for different values of reward probability for *N* = 6 model. Overall, there are 62 subsets of meta-states for *N* = 6 model. For superior models, the bottleneck subset (blue) is always across the plastic boundary to minimize noise for a given level of adaptability, whereas the rapidly mixing subsets (red) consist of only transient meta-states and thus build a quick connection between reservoir and buffer meta-states. (**d**) The APT in the plastic model and superior metaplastic models with different numbers of meta-states using Monte Carlo simulations.

The adaptability in the model depends on the rates of transitions among all subsets of meta-states, whereas noise (in reward estimation) depends on the flow across the plastic boundary (i.e. transitions between weak and strong meta-states and vice versa). Therefore, to understand how the model’s adaptability is optimized for a given level of noise, we computed the ‘effective transition rate’ for all subsets of meta-states. The effective transition rate was defined as the outward flow of synapses out of that subset, divided by the fraction of synapses in that subset, and is closely related to the concept of conductance in Markov chains (Sinclair and Jerrum, 1989) (Fig. 3a; see Methods). Importantly, the model’s adaptability is constrained by its slowest effective transition rate. In superior models, to reduce noise with a minimum cost to the adaptability, the slowest transition rates should be at the plastic boundary. We found that this was the case for all superior models (Fig. 2c). Interestingly, having the minimum effective transition rates at plastic transitions created a ‘bottleneck’ for the flow between weak and strong meta-states. This bottleneck helped reduce noise without significantly reducing the adaptability. The superior models with *N* > 4 also contained transient meta-states, with the fastest effective transition rates between buffers and reservoirs, resulting in improved adaptability (Fig. 2b-c and Fig. 3).

**Figure 3.**
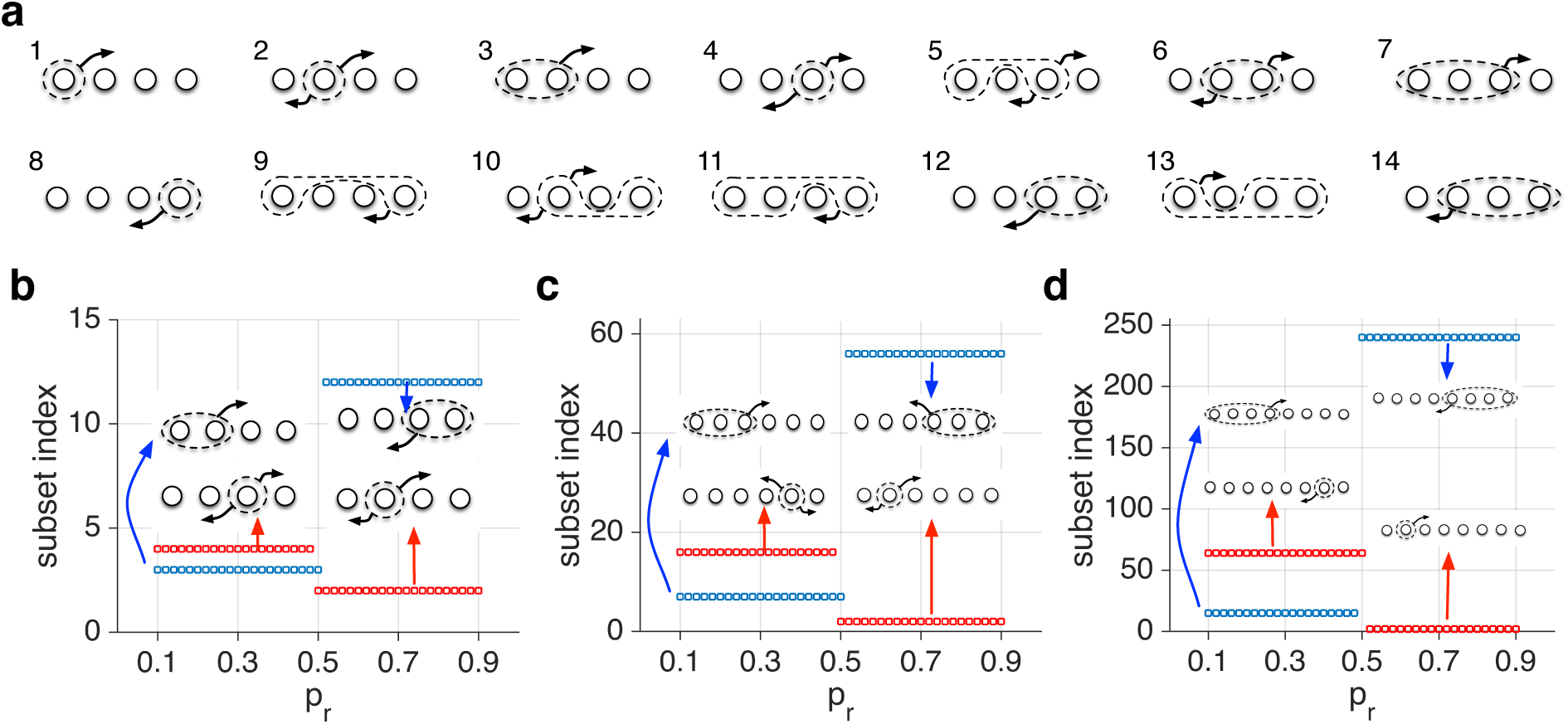
(**a**) Schematic of possible subsets of meta-states for *N* = 4 model. Overall, there are (2^*N*^ − 2) subsets of meta-states for a given metaplastic model. (**b-d**) Plotted is a subset of meta-states with the fastest and slowest effective transition rates for different values of reward probability for the metaplastic models with *N* = 4, 6, 8. For superior models, the bottleneck subset (blue) is always across the plastic boundary to minimize noise for a given level of adaptability, whereas the rapidly mixing subsets (red) consist of only transient meta-states in order to a build a quick connection between reservoir to buffer meta-states. Insets show the corresponding subset of meta-states.

This specific arrangement of meta-states and transitions between them, as well as the adjustment of the metaplastic model to reward probability, enabled metaplastic models to be more adaptable than corresponding plastic models. To demonstrate this superior adaptability, we used the effective learning rates for a given value of *p*_*r*_ to define an equivalent *N* = 2 for any metaplastic model. We found that metaplastic models showed larger sensitivity to reward probability than equivalent plastic models (Fig. 4). Moreover, metaplastic models were more adaptable and more precise than their equivalent plastic models. These results demonstrate that the dynamic adjustment of learning in metaplastic models is crucial for improving the APT, and that this adjustment cannot be achieved by simply exchanging the learning rates in corresponding plastic models for similar learning rates.

**Figure 4.**
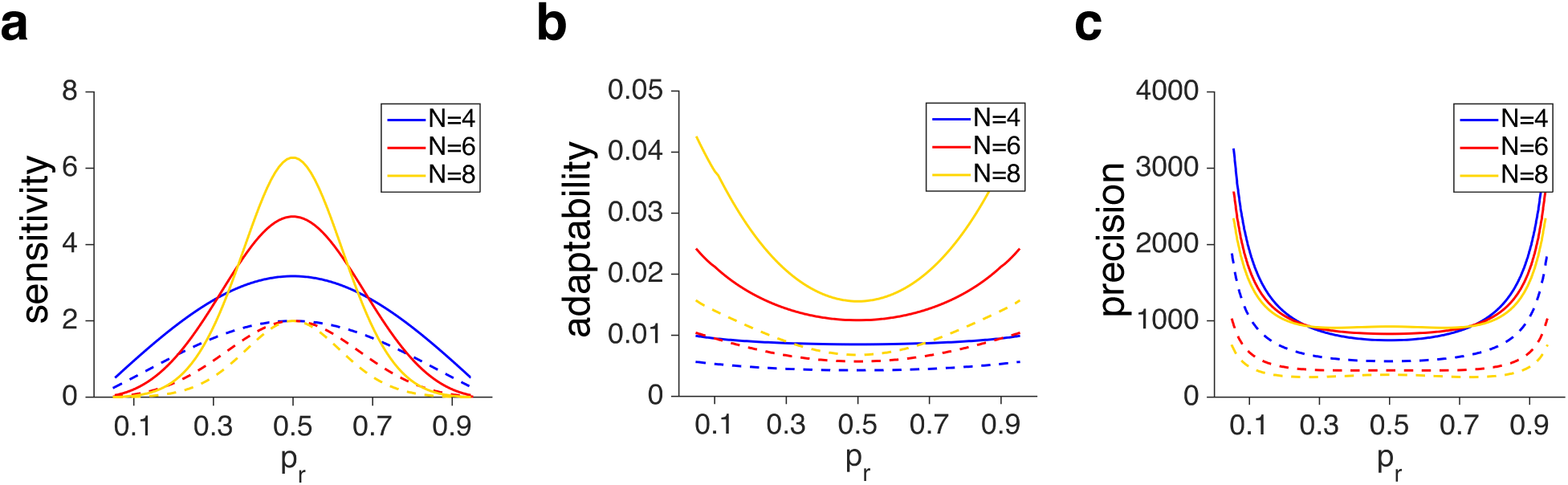
Comparisons between the behavior of the metaplastic models and equivalent plastic models with the effective learning rates for a given value of *p*_*r*_. (**a**) Plotted is sensitivity in three superior metaplastic models (solid curves) and their equivalent plastic models (dashed curves) as a function of *p*_*r*_. The equivalent plastic models are constructed using the effective learning rates for a given value of *p*_*r*_ and a metaplastic model. (**b-c**) Plotted is the adaptability and precision as a function of *p*_*r*_ for the same models presented in (a). The metaplastic models outperform equivalent plastic models in terms of sensitivity, precision, and adaptability for all values of reward probability.

To further study the characteristics of superior metaplastic models, we next examined the transition probabilities in these models. We found that most transition probabilities were very close to zero, allowing for the creation of reservoirs and buffers, while non-zero transition probabilities varied proportionally to create models with different levels of adaptability and precision. For example, in metaplastic models with four meta-states (*N* = 4), three of six transition probabilities for potentiation were zero, two others were equal, and the last one was very close to those two other non-zero probabilities (Supplementary Figure 3). Based on these observations, we constructed a simple family of metaplastic models using a single parameter. Even such simple metaplastic models can overcome the APT, and this ability was improved with additional meta-states (Fig. 5). Overall, these results show that metaplastic models outperform plastic models, not because they have more parameters, but because they have a structure that allows for adjustment of learning.

**Figure 5.**
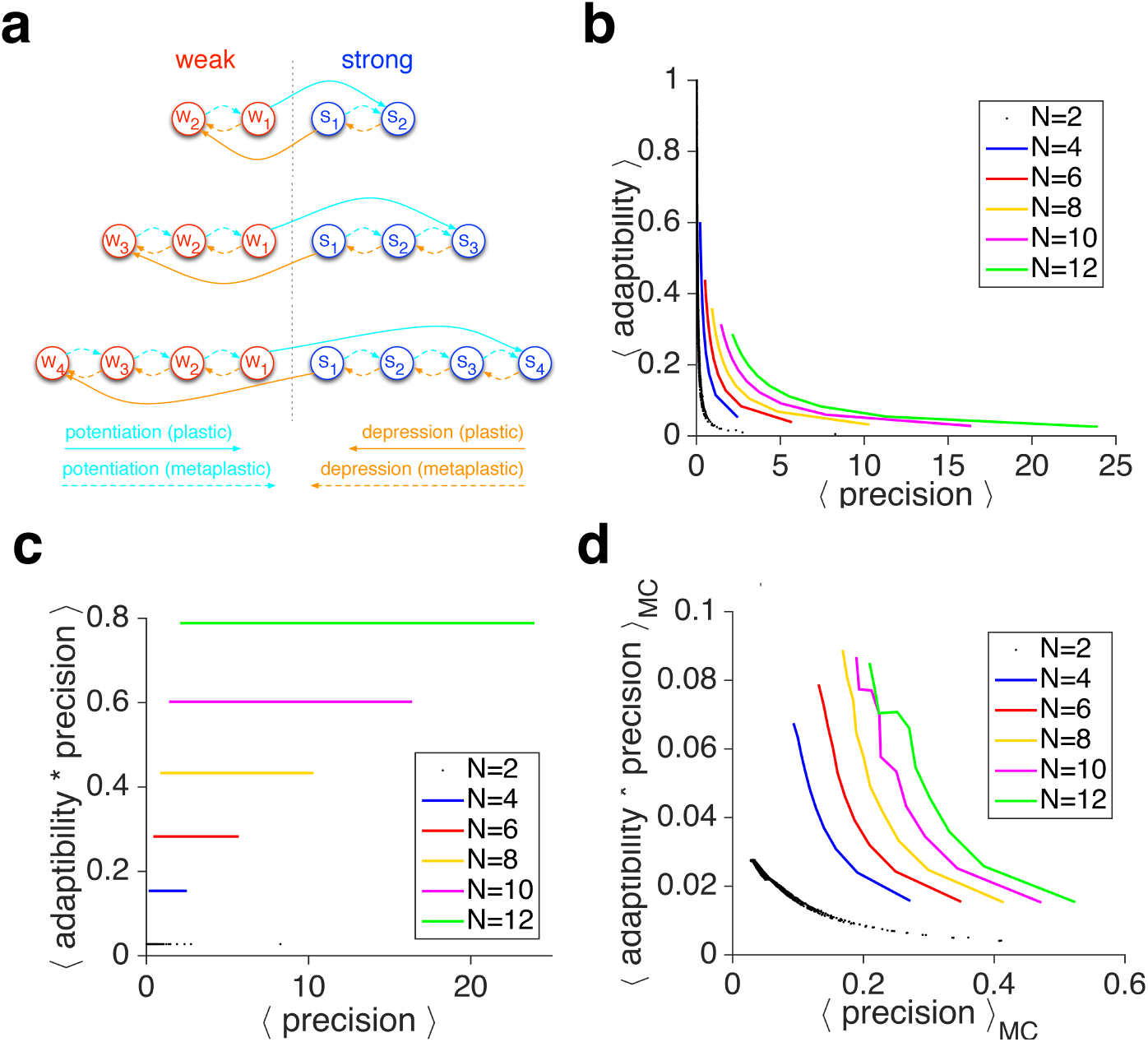
The APT in a special family of metaplastic models with only one parameter. (**a**) The structure of the special family of metaplastic models with only one parameter (for *N* = 4, 6, 8 meta-states). (**b**) The adaptability as a function of the precision for the simple metaplastic models using the mean-field approach. (**c**) The 𝔸 × 𝕡 as a function of the precision using the mean-field approach. (**d**) The same as in (c) but using Monte Carlo simulations. Increasing the number of meta-states improves the APT.

The results from one-parameter models also illustrate that having more meta-states can improve the ability of metaplasticity to overcome the APT. Nevertheless, the basic mechanism for this improvement is the existence of reservoirs, buffers, and a bottleneck for changing synaptic efficacy. Additional meta-states provide intermediate transitions between reservoirs and buffers that could increase signal and reduce noise without significantly decreasing the adaptability (Fig. 5). As a result, models with larger numbers of intermediate meta-states show better matching of sensitivity to noise as well as more optimized adaptability for a given level of noise. Essentially, the specific structure for changing synaptic efficacy allows the models with a large number of meta-states to collect evidence (by transitioning synapses to shallower meta-states) before making a change.

Even though the results above were obtained using a mean-field (MF) approach, our findings also hold using Monte Carlo (MC) simulations (Fig. 2d). The only difference between this approach and MC simulations was the estimation of noise, since the MF approach provides an accurate estimate of the signal and adaptability. Using MF, the estimated noise was set to one-step noise, which is equal to the weighted average of changes in the steady state of synaptic strength due to a potentiation and depression event. The one-step noise converges to the actual noise if the adaptability is equal to 1. When the adaptability is different from 1, one-step noise underestimates the actual level of noise measured by real simulations (Supplementary Figure 4a). Intuitively, this underestimation occurs because for MC simulations, fractions of synapses in different meta-states fluctuate around their steady-state values and thus add extra noise. While underestimation of noise increases with the number of meta-states, this effect is not strong enough to change the order of models in their ability to overcome the APT (compare Figures 2d with 1b) or the sensitivity-to-noise matching (Supplementary Figure 4b).

The ultimate test for whether metaplastic transitions are crucial for mitigating the APT is to replace these transitions with plastic ones (transitions that change synaptic efficacy) while keeping the same number of states and transitions. Therefore, we examined the APT in the simple family of metaplastic models, but with different values of synaptic efficacy assigned to different meta-states (Fig. 6a; see Methods). This ‘graded-plasticity’ model could be reduced to the metaplastic model by setting equal values of synaptic efficacy for different weak or strong states. We found that 𝔸 × 𝕡 monotonically increased as the graded-plasticity model became more similar to the metaplastic model (Fig. 6b-d). These results demonstrate that metaplastic transitions are crucial for mitigating the APT.

**Figure 6.**
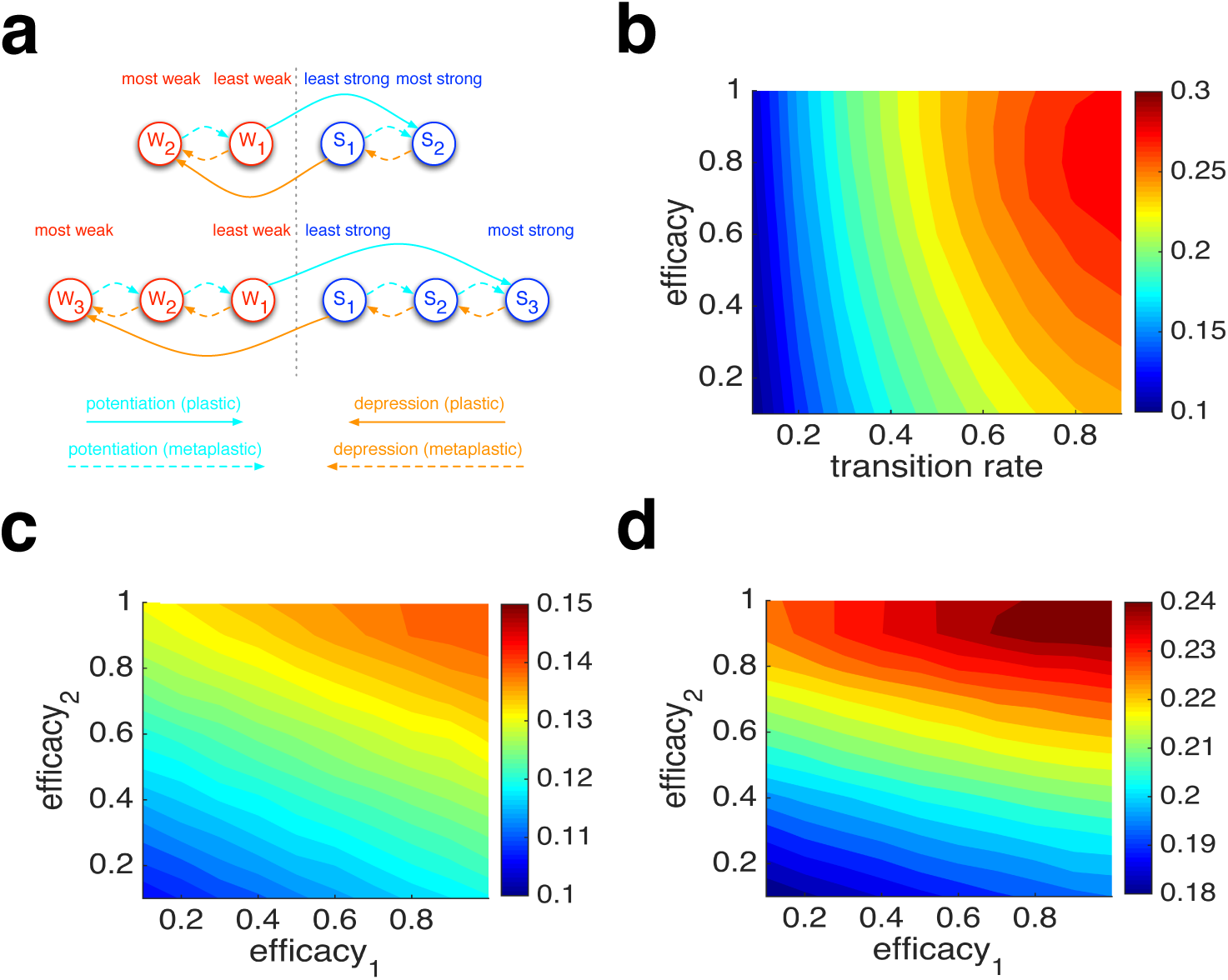
Graded-plasticity reduces the ability to overcome the APT, indicating that metaplasticity is crucial for adaptive learning. (**a**) Schematic of the simple-family graded-plasticity model. This model has an equal number of states and transitions as the metaplastic model, but with different values of synaptic efficacy assigned to different states. (**b**) Plotted is the average 𝔸 × 𝕡 in the graded-plasticity model with four states (*N* = 4), as a function of the single transition rate and the efficacy of the least weak state (*efficacy* = 1 is equivalent to the *N* = 4 meta-plastic model). (**c**) Plotted is the average 𝔸 × 𝕡 in the graded-plasticity model with six states (*N* = 6), as a function of the efficacy of the weaker states when the single transition rate was set to 0.2 (*efficacy*_*1*_ = *efficacy*_*2*_ = 1 is equivalent to the *N* = 6meta-plastic model). (**d**) The same as in (c) but for the transition rate equal to 0.7.Overall, 𝔸 × 𝕡 monotonically increased as the graded-plasticity model became similar to the metaplastic model.

## Discussion

The demands of learning in a changing world require a high degree of adaptability, which comes at the cost of low precision (Farashahi et al., 2017b). Here we show how metaplasticity, which is reflected in the unreliability of synaptic plasticity, can provide a solution for substantially overcoming the APT. More specifically, by optimizing the APT for a given level of precision, we identify non-trivial characteristics of superior metaplastic models. The superior models contain reservoir and buffer meta-states; synapses in reservoir meta-states do not change their efficacy upon reward feedback, whereas those in buffer meta-states can change their efficacy. Moreover, rapid changes in efficacy are limited to synapses occupying buffers, which provides a bottleneck that reduces noise without significantly decreasing adaptability. In contrast, more-populated reservoirs can generate a strong signal without manifesting any observable plasticity. The generation of reservoirs and buffers by metaplastic synapses results in the adjustments of learning, or the degree of plasticity, according to recent reward history. For example, when synapses occupy reservoir meta-states, which occurs with consecutive rewarded or unrewarded trials in a stable environment, the behavior should become less adaptable. However, when reward history changes over time, synapses mainly occupy buffer meta-states, causing more adaptable behavior. Overall, the model predicts that learning should be more sensitive to the reward sequence than what has previously been assumed.

Importantly, the results of one-parameter models and MC simulations show that having more meta-states can improve the metaplasticity for overcoming the APT and, in addition, gives rise to more robust models for adaptive learning. The basic mechanism for this improvement is the generation of reservoirs and buffers that create a bottleneck for changing synaptic efficacy; additional meta-states provide intermediate transitions between reservoirs and buffers that could reduce noise without compromising adaptability. Interestingly, it has been shown that in the framework of Markov chains, the eigenvalues and eigenvectors of models with bigger spectral gaps (i.e. more adaptable) are less sensitive to perturbation of transition probabilities (Funderlic and Meyer 1986; Seneta 1993; Meyer 1994; Cho and Meyer 2001). In other words, more-adaptable models can produce signals without fine-tuning. Superior metaplastic models require only a few parameters, and their behavior is not very sensitive to these parameters.

As a higher-order form of plasticity, metaplasticity has been successfully used to explain paradoxical observations regarding synaptic plasticity by considering prior synaptic activity (Yger and Gilson 2015). The computational power of metaplastic synapses has only recently been explored to address memory retention (Fusi et al., 2005; Fusi and Abbott, 2008; Barrett and van Rossum, 2009; Lahiri and Ganguli, 2013; Farashahi et al., 2017a), but its benefit for reward-dependent learning remains unknown. Our results can be applied to estimating signals other than reward probability and can be generalized to other domains of learning. For example, it has been shown that a tradeoff between adaptation speed and accuracy is limited by the power dissipation because adaptation processes are necessarily dissipative (Lan et al., 2012). We found that in superior models, many transitions can occur between meta-states without any changes in efficacy. Considering that changing efficacy is energetically costly, this finding may suggest the importance of energy constraint for neural computations underlying learning (Laughlin et al., 1998; Lennie, 2003).

Our results could also explain why plasticity protocols are unreliable. As we showed, superior metaplastic models create bottlenecks for changing synaptic efficacy since such a property can reduce noise without sacrificing adaptability. However, having plastic transitions to be limited to those that occur from buffers would make many transitions invisible to measurement of change in synaptic efficacy. Therefore, until such a structure is specifically tested, plasticity protocols will be perceived as noisy and unreliable.

Our proposal provides a new approach for studying synaptic plasticity and its contribution to brain computations. Our model predicts that a previous reward outcome (learning experience) not only contributes to learning and behavioral changes, but also affects subsequent induction of such changes within a specific time window. On the one hand, certain sequences of reward feedback cause the nervous system to become more receptive to subsequent similar feedback. On the other hand, consecutive feedback can shape future learning such that it is not responsive to feedback in the opposite direction. Understanding such propensity and unresponsiveness to reward feedback could provide new insights into habit and addiction, respectively. Therefore, further investigations into metaplasticity, both at the behavioral and synaptic levels, could help researchers find tools for improving learning, especially as regards habits and addiction (Moussawi et al 2009; Hulme et al., 2013).

## Methods

### Metaplastic model

Our general model of metaplasticity consisted of multiple meta-states associated with two values of synaptic efficacy (weak and strong), and all possible transitions between these meta-states (Fig. 1a). The metaplastic models have *N* distinct meta-states, half of which are associated with strong synaptic efficacy and half with weak. The model is completely specified with two transition matrices, one for a potentiation event 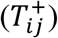 and one for a depression event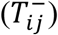, corresponding to rewarded and unrewarded trials, respectively. Here, we assumed that metaplastic transitions have a consistent order such that potentiation and depression events (on rewarded and unrewarded trials, respectively) create flows in opposite directions. This assumption also establishes weak and strong meta-states with different ‘depths’ such that deeper states are further from the plastic boundary (Fig. 1a). Moreover, we assumed symmetry between information by reward and no-reward feedback, and thus only focused on mirror-symmetric flows. This assumption put another constraint on the potentiation and depression matrices:

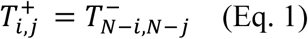

Based on these assumptions, transition matrices for potentiation and depression events can be represented by lower-triangular and upper-triangular matrices:

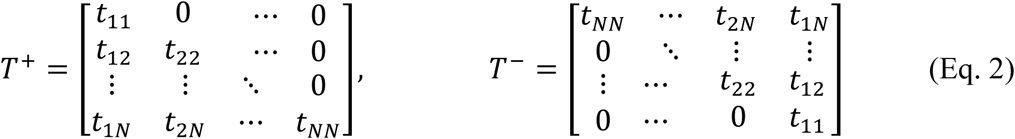

There are *N*(*N* − 1)/2 unique transition probabilities for models with *N* meta-states. The probability conservation was dictated by the transition flows out of any meta-state summing up to 1.

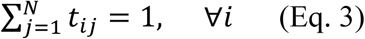

### Mean-Field approach

At any point in time, the signal (*S*) was defined as the difference between the fractions of synapses in the strong and weak meta-states,

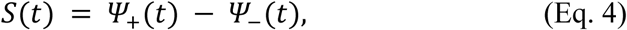

where 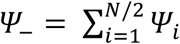 and 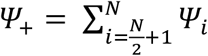 are fractions of synapses in the weak and strong meta-states, respectively. In the mean-field (MF) approximation approach, the average system dynamics is fully described by the average transition matrix for a given reward probability 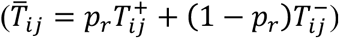. The eigenvector, *Ψ*, with an eigenvalue *λ* = 1 (the largest eigenvalue according to Perron-Frobenius theorem) of average transition matrix, 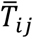, provided the steady state of the model from which the average signal was calculated using Equation 4.

As a proxy for signal fluctuations around its average value, we introduced the concept of ‘one-step noise’ as the mean magnitude deviation from the average signal due to one potentiation or depression event:

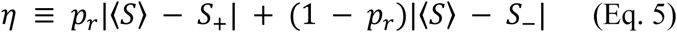

where *S* is the average signal based on the steady-state solution, and *S*_+_ and *S*_−_ are the signal values after the application of the potentiation or depression transition matrices on the steady-state solution, respectively. In general, noise at time (*t* + 1) is a combination of several components: (1) the attenuated transferred noise from the state of the system at time *t*; (2) the amount of noise generated in one step, from *t* to (*t* + 1); (3) the inherent noise involved in translating *p*(*t*) to a binary representation with potentiation and depression events; and finally (4) a finite size effect when dealing with a limited number of identical synapses. The one-step noise measures the second component and always underestimates the level of the noise in our system. In contrast, the Monte Carlo simulation contains the sum of the first three components mentioned above.

The Monte Carlo simulations were performed by running multiple trials starting from a given initial state in environments with identical reward statistics (reward probability was the same but the reward sequence varied across different simulations). Data from an initial relaxation period was thrown away in order to remove dependence on the initial state, and the relevant quantities were computed by averaging over the ensemble at a given time step or across time. Moreover, to further reduce the relaxation time, we started from the steady-state solution of the mean-field equation for the initial environment.

We defined precision as the ratio of the signal sensitivity and the one-step noise:

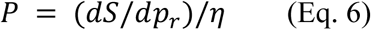

Therefore, precision measures the discriminability between two adjacent reward probabilities based on their resulting signals. We chose this measure instead of the difference between the estimated and actual reward probability because the firing rate of neurons, which represent reward values, can be differentially scaled by their input firing rates, and so this difference is irrelevant.

Finally, the adaptability of the model was defined as the rate of the decaying mode in the system, and was estimated using the difference between the second-largest eigenvalues (slowest decaying mode) of the average transition matrix and 1 (𝔸 = 1 − *λ* _2_), also known as the spectral gap in the Markov chains literature. Therefore, adaptability measures the lower bound for the speed of the system in converging to its final steady state.

By focusing on the steady-state solution, the concept of learning rates in the plastic models (*N* = 2) can be generalized to higher *N* as the effective learning rates,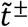. The effective learning rates were defined as the relative change in the fraction of synapses in the weak or strong meta-states after a potentiation or depression event:

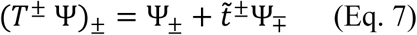

where ()_±_ is the sum of the fraction of strong/weak meta-states. To examine transitions from a given subset of meta-states, we also defined the ‘effective transition rate’ as the outward flow of synapses from that subset, divided by the fraction of synapses in that subset (Fig. 3a). The effective transition rate 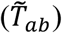 assigns a single rate for outward transition from a set of meta-states *a* to a set of meta-states *b*. There are (2^*N*^ − 2) non-trivial ways that *N* meta-states can be partitioned into two disjoint, complementary subsets.

A closely related concept of conductance, *C*(*S*), for a given subset *S* in a Markov chain is defined as the outward flow from that subset divided by the minimum of occupancy in that subset, *π*(*S*), and occupancy in its complementary set *π*(*S*^*c*^). The magnitude of one-step noise is directly related to the effective transition rate when the two subsets are chosen based on their synaptic efficacy. The value of spectral gap (i.e. the difference between the second-largest eigenvalues of the average transition matrix and 1) is constrained by the minimum conductance among all possible subsets of states (Sinclair and Jerrum, 1989).

### Finding the optimized solutions

Finding optimized solutions (i.e. upper-boundary in adaptability × precision vs. precision plot) was performed in two stages. An initial upper envelope in the 𝔸 × 𝕡 vs. 𝕡 (using discretization for 𝕡) was constructed by random sampling of 10^7^ transition matrices. The transition matrices were divided into *n* bins according to their precision, 𝕡, and the transition matrix with the highest value of 𝔸 × 𝕡 in each bin was selected. These transition matrices were then used as the initial points for our optimization process. To avoid local minima, at the beginning of each iteration, a duplicated copy of the initial transition matrix with added small jitters was generated. All the resulting 2*n* transition matrices were used as the starting point of our optimization. At the end of each optimization iteration, the best solutions in each bin were selected out of all initial transition matrices and the final outcome of our optimization procedure and used for the initial samples of the next iteration. For high-dimensional cases (*N* > 4), we went through multiple iterations of the optimization process. The higher dimensional solutions are more robust against fluctuations, and optimized solutions can be found by increasing the bin numbers (initial points) and the number of optimization iterations. The optimization was constrained by keeping the sum of every column in transition matrices with positive elements to one. We used MATLAB’s ‘fminsearch’ function for the basic optimization process.

## Supplemental Information

Supplemental Information includes 1 note and 4 figures.

## Author contributions

P.K. and A.S. conceived the model and designed the research. P.K. performed model simulations and analyzed the data. P.K. and A.S. interpreted simulation results and wrote the manuscript.

## Acknowledgments

We would like to thank Brad Duchaine, Daeyeol Lee, and Matt van der Meer for helpful comments on the manuscript. This work was supported by startup fund from Dartmouth College and the Neukom Institute CompX Grant to A.S.

## Supplementary Materials

### Supplementary Note 1: Adaptability−precision tradeoff in RL and model with plastic synapses

#### Equivalence of the RL model to the model with plastic synapses

Here we show that stochastic synapses without metaplasticity are equivalent to the RL model based on reward prediction error (RPE). A standard RL model based on reward prediction error (the difference between expected and actual outcomes) and with equal learning rates for rewarded and unrewarded trials can estimate the reward probability (Sutton and Barto, 1998). This RL model is fully described by its value function, *V*. Its temporal dynamics in response to a feedback sequence is governed by a learning rate *δ* and the reward prediction error which is the difference between expected and actual reward.

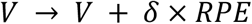

where *RPE* = (1 − *V*) or (−*V*) on rewarded or unrewarded trials, respectively. The learning rates could be different on rewarded and unrewarded trials (*δ*_+_, *δ*_−_) resulting in the following update rules:

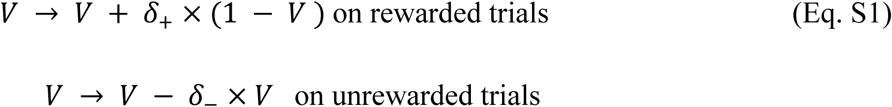

The average of *V* approaches the reward probability *p*_*r*_ in the environment when *δ*_+_ = *δ*_−_.

A model with binary synapses (‘weak’ and ‘strong’ states) that undergo stochastic reward-dependent plasticity can also provide an unbiased estimate of the reward probability (Soltani and Wang, 2006; Soltani et al., 2006). In this model, weak synapses can be potentiated on rewarded trials with a probability *t*^+^ (potentiation rate), whereas strong synapses can be depressed on unrewarded trials with a probability *t*^-^ (depression rate). Therefore, the fraction of synapses in the strong state, Ψ_+_, is updated as the following:

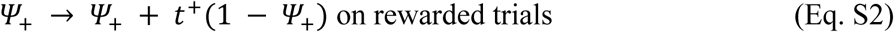

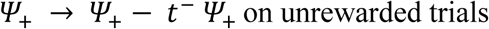

The equivalence between the RL and plastic models (*N* = 2) can be seen by comparing the strong-state occupancy percentage with the value function and replacing *δ* _±_ with 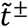 in Equations S1 and S2.

In the model with plastic synapses (*N* = 2), we defined the signal as the difference between the fractions of synapses in the strong and weak states, *S* = *Ψ*_+_ − *Ψ*_−_. When the reward probability is equal to *p*_*r*_, the average signal is equal to

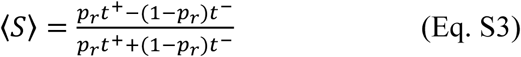

Therefore, the signal ‘sensitivity’, defined as the derivative of the average signal with respect to *p*_*r*_, is equal to

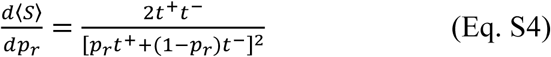

The ‘one-step noise’, defined as the mean magnitude deviation from the average signal in one time step, is equal to

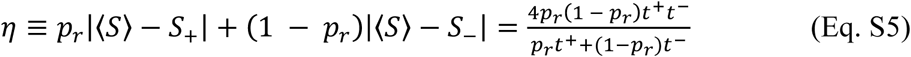

Hence, the precision is equal to

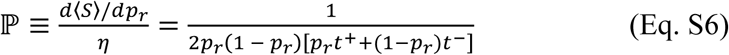

The adaptability is defined as the rate of decaying mode in the system. In *N* = 2 models, the rate of approach toward the average signal is fully governed by the weighted average of the learning rates:

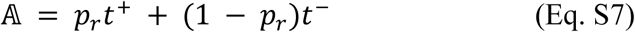

Finally, *N* = 2 models show a strict APT since the product of adaptability and precision is independent of model parameters and only depends on reward probability:

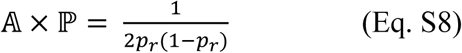

In general, adaptability and noise are related to each other in the model with plastic synapses and in the RL model. This is because an increase in transition probabilities between weak and strong states causes larger flows between the two states, which in turn increases noise. Importantly, adopting different transition probabilities (or learning rates in RL) for potentiation and depression events cannot improve the APT, rather affects the average values for adaptability and precision individually (Fig. 1b and Supplementary Figure 1).

**Figure S1.**
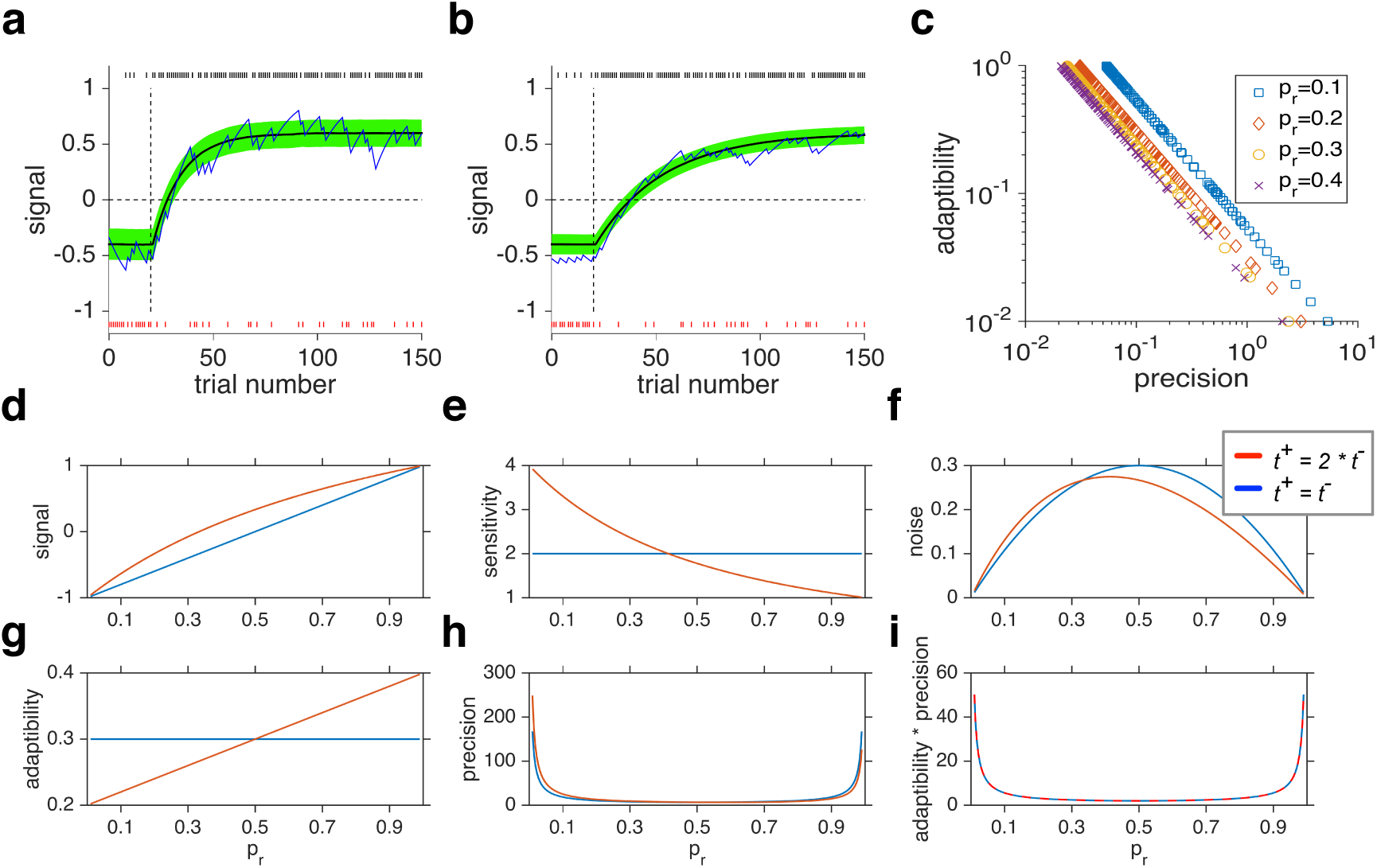
The APT in the plastic model (*N* = 2). (**a-b**) Dynamic of the signal in the plastic model in response to a sudden change in reward probability. Decreasing the transition rates (from *t*^+^ = *t*^−^ = 0.07 in (a) to *t*^+^ = *t*^−^ = 0.03 in (b)) resulted in noise reduction in the asymptotic value of the signal, but at the expense of slower convergence to this asymptotic value. (**c**) The APT manifests itself for different learning rates for different reward probabilities. Plotted is the adaptability, as a function of the precision for different values of *p*_*r*_. Each dot corresponds to a specific set of parameter values. The APT is stronger as *p*_*r*_ becomes closer to 0.05 (see Equation S8 in the Supplementary Note 1). (**d-i**) Characteristics of the plastic model, measured using different quantities, as a function of reward probability for two sets of learning rates (*t*^+^ = 2 × *t*^−^ = 0.4 and *t*^+^ = 2 × *t*^−^ = 0.3). Adopting different learning rates improves the adaptability for certain values of *p*_*r*_ and improves precision for complementary of values of *p*_*r*_, resulting in a strict tradeoff between adaptability and precision. A simple RL model based on RPE behaves similarly to the plastic model shown here.

**Figure S2.**
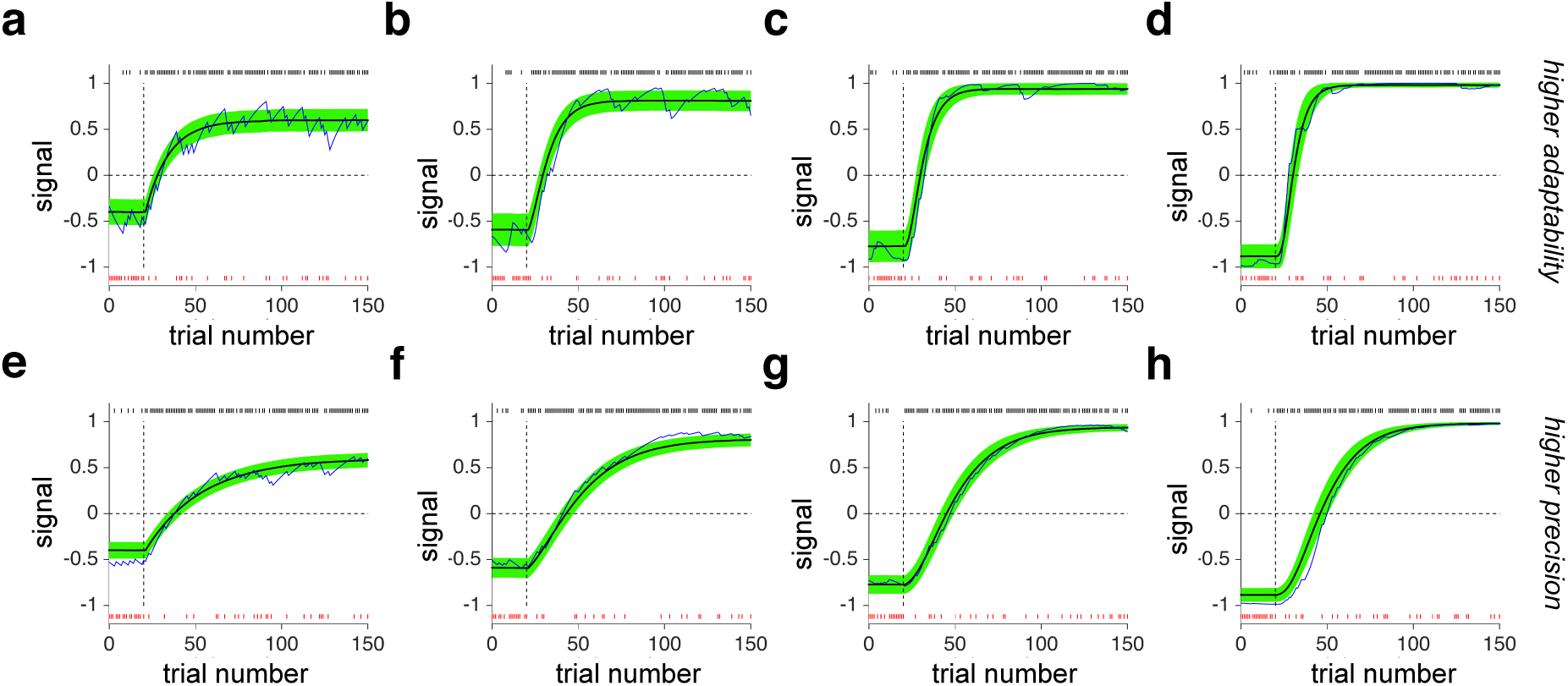
Example traces of the signal in response to a sudden change in reward probability in the plastic model (*N* = 2), and three superior metaplastic models with different numbers of meta-states. In each plot, the blue trace is an example estimate based on the shown reward sequence (tick marks on the top and bottom correspond to rewarded and unrewarded trials, respectively). The reward probability changed from 0.3 to 0.8 on trial 24. The black curve shows the average signal, and the green shade shows the signal plus/minus its s.e.m. Models in (a-d) are more adaptable, whereas models in (e-h) are more precise. For these simulations, example models were selected to have the same average precision. Overall, metaplastic models can improve the adaptability without increasing noise in the signal (thinner green lines).

**Figure S3.**
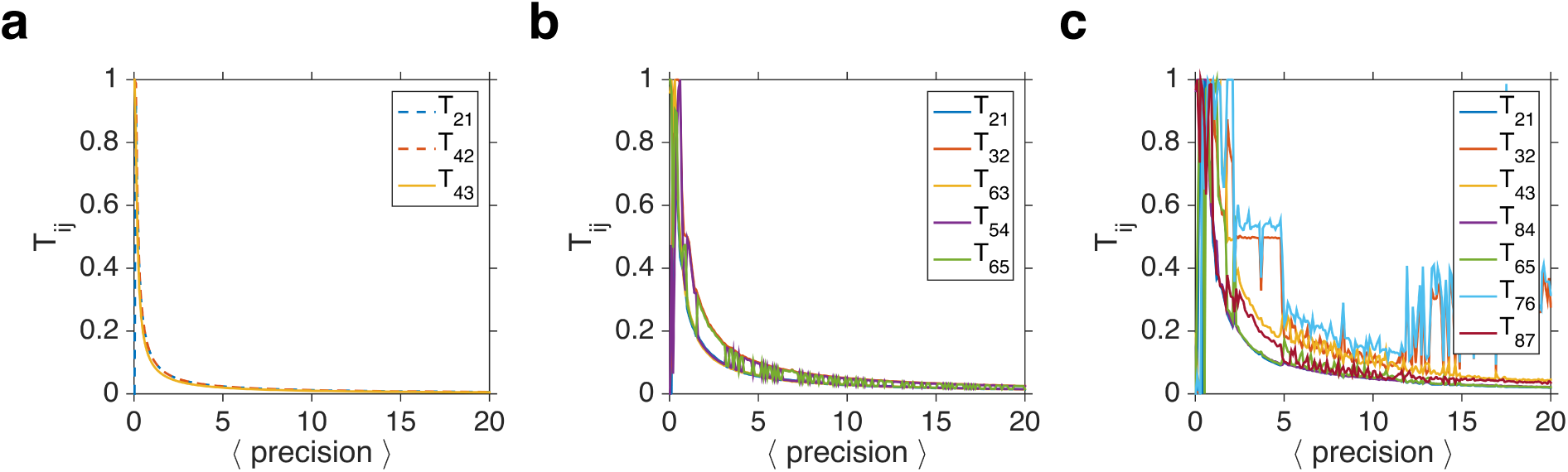
Transition probabilities in the superior metaplastic models with different numbers of meta-states (*N* = 4, 6, 8). (**a-c**) Plotted are the transition probabilities for superior models for a given value of average precision. Only a few transition probabilities are non-zero, and the rest vary together, revealing the specific structure of metaplasticity that is useful for overcoming the APT. For models with larger numbers of meta-states, finding superior models is more difficult because those models are more robust against fluctuations in the model parameters.

**Figure S4.**
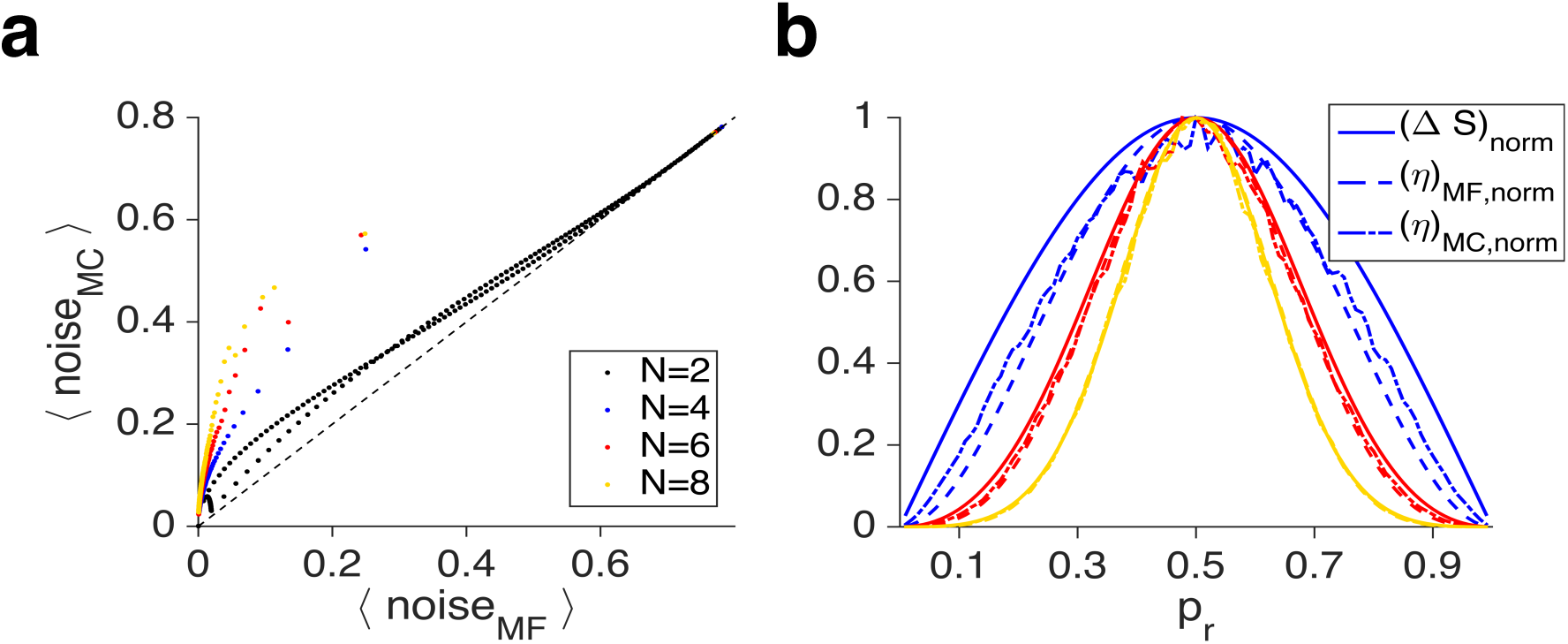
Comparison of one-step noise and noise based on Monte Carlo simulations. (**a**) Plotted is the noise computed using Monte Carlo simulations as a function of one-step noise for the plastic model and superior metaplastic models with different numbers of meta-states. One-step noise sets a lower bound for simulation noise. The mean-field approximation for noise becomes more accurate for higher adaptability. (**b**) Comparison of sensitivity-to-noise matching based on one-step noise and Monte Carlo simulations.

